# ZC3H4 restricts non-coding transcription in human cells

**DOI:** 10.1101/2021.02.10.430684

**Authors:** Chris Estell, Lee Davidson, Pieter C. Steketee, Adam Monier, Steven West

## Abstract

The human genome encodes thousands of non-coding RNAs. Many of these terminate early and are then rapidly degraded, but how their transcription is restricted is poorly understood. In a screen for protein-coding gene termination factors, we identified ZC3H4. However, its depletion causes upregulation and extension of hundreds of unstable transcripts, particularly antisense RNAs and those transcribed from so-called super-enhancers. These loci are occupied by ZC3H4, suggesting that it directly functions in their transcription. Consistently, engineered tethering of ZC3H4 to reporter RNA promotes its degradation by the exosome. ZC3H4 is metazoan-specific - interesting when considering its impact on enhancer RNAs that are less prominent in single-celled organisms. Finally, ZC3H4 loss causes a substantial reduction in cell proliferation, highlighting its overall importance. In summary, we identify ZC3H4 as a factor that plays an important role in restricting non-coding transcription in multi-cellular organisms.

## INTRODUCTION

Most of the human genome can be transcribed by RNA polymerase II (Pol II). Among these transcripts are thousands of long non-coding RNAs, broadly classified as greater than ~200 nucleotides in length (Kopp and Mendell, 2018). They share some structural features with coding transcripts, but most of them are rapidly degraded by the exosome (Davidson et al., 2019; Preker et al., 2008; Schlackow et al., 2017). Their degradation is coincident with or shortly after transcriptional termination, which often occurs within 1-2 kilobases (kb). The mechanisms for terminating non-coding transcription are poorly understood, especially by comparison with those operating at protein-coding genes.

Termination of protein-coding transcription is coupled to 3’ end processing of pre-mRNA via cleavage at the polyadenylation signal (PAS) (Eaton and West, 2020). A PAS consists of an AAUAAA hexamer followed by a U/GU-rich region (Proudfoot, 2011). After assembly of a multi-protein processing complex, CPSF73 cleaves the nascent RNA and the Pol II-associated product is degraded 5’→3’ by XRN2 to promote termination (Eaton et al., 2018; Eaton et al., 2020; Fong et al., 2015). The Pol II elongation complex is modified as it crosses the PAS, which facilitates its termination by XRN2 (Cortazar et al., 2019; Eaton et al., 2020). Depletion of XRN2 or CPSF73 causes read-through downstream of some long non-coding genes (Eaton et al., 2020). However, a substantial fraction of non-coding transcription is insensitive to their depletion suggesting the use of alternative mechanisms.

The Integrator complex aids termination of many non-coding transcripts, with the archetypal example being snRNAs (Baillat et al., 2005; Davidson et al., 2020; O’Reilly et al., 2014). The mechanism is analogous to that at protein-coding genes, driven by endonucleolytic cleavage that is supplied by INTS11. However, INTS11 activity does not precede XRN2 effects (Eaton et al., 2018). Moreover, while CPSF73 is indispensable for termination at protein-coding genes, a milder effect is seen at snRNAs when Integrator is depleted (Davidson et al., 2020). Integrator is also implicated in the termination of promoter upstream transcripts (PROMPTs) and enhancer RNAs (eRNAs) (Beckedorff et al., 2020; Lai et al., 2015; Nojima et al., 2018). Since relatively mild termination defects are associated with the absence of these factors, there may be additional mechanisms. Indeed, CPSF and the cap binding complex-associated factor, ARS2, are both implicated in the termination of promoter-proximal transcription (Iasillo et al., 2017; Nojima et al., 2015).

A variety of processes attenuate transcription at protein-coding genes (Kamieniarz-Gdula and Proudfoot, 2019). Frequently, this is via premature cleavage and polyadenylation (PCPA) that can be controlled by U1 snRNA, CDK12, SCAF4/8 or PCF11 (Dubbury et al., 2018; Gregersen et al., 2019; Kaida et al., 2010; Kamieniarz-Gdula et al., 2019). PCPA is frequent on many genes since acute depletion of the exosome stabilises its predicted products in otherwise unmodified cells (Chiu et al., 2018; Davidson et al., 2019). Integrator activity also attenuates transcription at hundreds of protein-coding genes, which is sometimes reversed by its absence leading to more full-length transcription (Elrod et al., 2019; Lykke-Andersen et al., 2020; Tatomer et al., 2019).

A less-studied termination pathway at some intragenic non-coding regions is controlled by WDR82 and its associated factors (Austenaa et al., 2015). In mammals, WDR82 forms at least two complexes: one with the SETD1 histone methyl-transferase and another composed of protein-phosphatase 1 and its nuclear targeting subunit PNUTS (Lee et al., 2010; van Nuland et al., 2013). A version of the latter promotes transcriptional termination in trypanosomes (Kieft et al., 2020) and the budding yeast homologue of WDR82, Swd2, forms part of the APT termination complex (Nedea et al., 2003). In mice, depletion of either WDR82, PNUTS or SET1 causes non-coding transcriptional termination defects. Notably, PNUTS/PP1 is implicated in the canonical termination pathway at protein-coding genes where its dephosphorylation of SPT5 causes deceleration of Pol II beyond the PAS (Cortazar et al., 2019; Eaton et al., 2020).

Here, we performed a proteomic screen for new termination factors by searching for proteins that bind to Pol II complexes in a manner that depends on PAS recognition by CPSF30. This uncovered ZC3H4, a metazoan zinc finger-containing factor without a characterised function in transcription. Although we anticipated a role for ZC3H4 in 3’ end formation, its effects on this process are mild and restricted to a small number of genes. Its main function is to restrict non-coding transcription, especially of PROMPT and eRNA transcripts, which are extended by hundreds of kb when ZC3H4 is depleted. ZC3H4 interacts with WDR82, the depletion of which causes similar defects. Tethered function assays show that ZC3H4 recruitment is sufficient to restrict transcription and cause RNA degradation by the exosome. In sum, we reveal ZC3H4 as a hitherto unknown terminator of promoter-proximal transcription with particular relevance at non-coding loci.

## RESULTS

### The effect of CPSF30 depletion on the Pol II-proximal proteome

The first step of PAS recognition involves the binding of CPSF30 to the AAUAAA signal (Chan et al., 2014; Clerici et al., 2018; Sun et al., 2018). We reasoned that elimination of CPSF30 would impede PAS-dependent remodelling of elongation complexes and cause the retention or exclusion of potentially undiscovered transcriptional termination factors. We used CRISPR/Cas9 genome editing to tag *CPSF30* with a mini auxin-inducible degron (mAID) (Figure 1A). The integration was performed in HCT116 cells where we had previously introduced the plant F-box gene, *TIR1*, required for the AID system to work (Eaton et al., 2018; Natsume et al., 2016). CPSF30-mAID is eliminated by 3 hours of indol-3-acetic acid (auxin/IAA) treatment (Figure 1B). This results in profound and general transcriptional read-through downstream of protein-coding genes (Figure 1C and S1A) demonstrating widespread impairment of PAS function.

**Figure 1:**
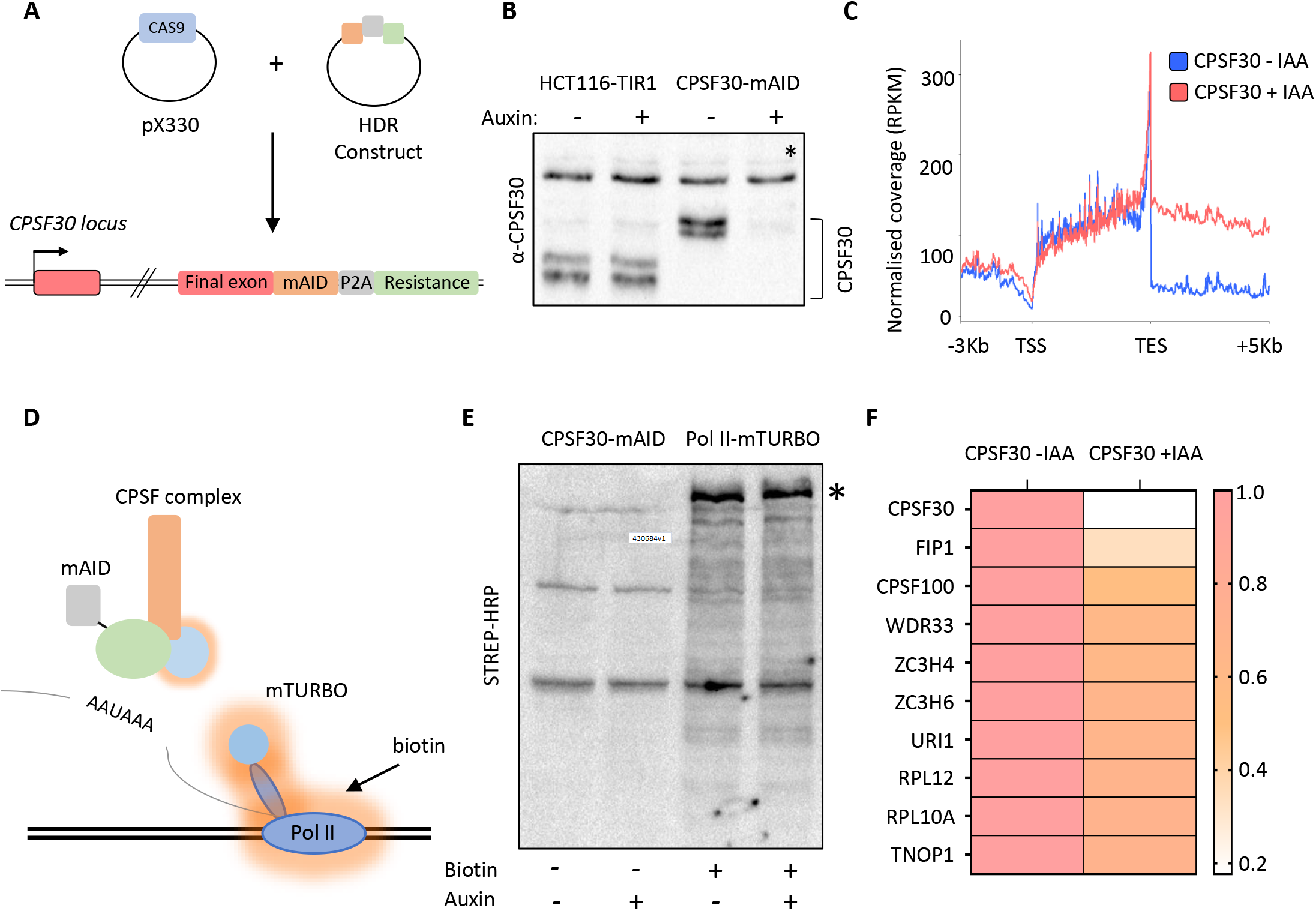
Proximity labelling of CPSF30-sensitive Pol II interactions by mTurbo. a) Schematic of strategy used to tag CPSF30 with the mini auxin inducible degron (mAID). Guide RNA-expressing Cas9 plasmid and homology directed repair (HDR) plasmids are shown and the resulting modification to *CPSF30* is represented with each inserted element labelled. b) Western blot demonstrating CPSF30 depletion. Parental HCT116-TIR1, or *CPSF30-mAID* cells, were treated +/− auxin for 3 hours then blotted. CPSF30 protein is indicated together with a non-specific product, marked by an asterisk, used as a proxy for protein loading. c) Metagene analysis of 1795 protein-coding genes demonstrating increased downstream transcription, derived from sequencing nuclear RNA, following auxin treatment (3hr) of *CPSF30-mAID* cells. TSS = transcription start site, TES = transcription end site, read through signal is normalised against gene body. RPKM is Reads Per Kilobase of transcript, per Million mapped reads. d) Schematic of our strategy to identify new factors involved in transcription termination. *CPSF30-mAID* cells edited to contain Rpb1-mTurbo (blue circle on Pol II). Addition of biotin induces mTurbo-mediated biotinylation (orange haze) of factors proximal to Pol II. CPSF complex is shown as an example of what might be captured by this experiment. e) Western blot showing streptavidin HRP probing of extracts from *CPSF30-mAID: RPB1-mTurbo* cells. Prior treatment with auxin (3hr)/biotin (2 mins) is indicated. The high molecular weight species in the + biotin samples corresponds in size to Rpb1-mTurbo (*). f) Heat map detailing proteins with the largest decrease in Pol II interaction. Data underpinning heat map are from mass spectrometry analysis of streptavidin sequestered peptides (+/− CPSF30) performed in triplicate. Labelling was for 2 minutes.

To identify Pol II interactions sensitive to CPSF30, we further modified *CPSF30-mAID* cells to homozygously tag the largest subunit of Pol II, Rpb1, with a mini(m)-Turbo tag (Figure 1D and S1B). mTurbo is an engineered ligase that biotinylates proximal proteins when cells are exposed to biotin (Branon et al., 2018). This occurs within minutes of biotin addition to culture media, which is advantageous for analysing dynamic proteins such as Pol II. We chose this approach rather than immunoprecipitation (IP) for two reasons: it allows isolation of weak/transient interactions (potentially disrupted during conventional IP) and may identify relevant proximal proteins that do not interact with Pol II directly. Importantly, CPSF30-mAID depletion still induced strong read-through in this cell line (Figure S1C).

*CPSF30-mAID:RPB1-mTurbo* cells were exposed to biotin before western blotting with streptavidin horseradish peroxidase (HRP). This revealed multiple bands with a prominent one corresponding in size to Rpb1-mTurbo and indicating the biotinylation of its proximal proteome (Figure 1E). A small number of bands were observed without biotin treatment due to endogenously biotinylated factors. Biotin-exposed samples were subject to tandem mass tagging (TMT) with mass spectrometry. We focused on proteins with reduced abundance after auxin treatment. The factor most depleted was CPSF30, confirming that its loss is reflected in the data (Figure 1F). As expected, Rpb1 was the most abundant factor in all samples consistent with its self-biotinylation seen by western blotting (Supplementary Excel file 1). After CPSF30, the most depleted factors were Fip1, CPSF100 and WDR33, which are in the CPSF complex. This implies that the major effect of CPSF30 depletion on the Pol II interactome is to prevent the recruitment/assembly of the CPSF complex. Otherwise, surprisingly few proteins showed reduced signal following auxin treatment.

### ZC3H4 is a putative termination factor that is metazoan-specific

ZC3H4 and ZC3H6 were the next most depleted. Both factors contain CCCH zinc finger motifs flanked by intrinsically disordered regions (Figure S2A). Their potential relationship to canonical 3’ end formation factors is suggested via the STRING database (Jensen et al., 2009) (Figure S2B). ZC3H4 is also co-regulated with mRNA processing factors suggesting a role in RNA biogenesis (Figure 2A; (Kustatscher et al., 2019)). Although little is reported on ZC3H4, two independent studies uncovered it as an interaction partner of WDR82 using Mass Spectrometry (Lee et al., 2010; van Nuland et al., 2013). WDR82 plays a key role in transcriptional termination in yeast, trypanosomes and mice (Austenaa et al., 2015; Kieft et al., 2020; Nedea et al., 2003). To verify this interaction, we tagged ZC3H4 with GFP and performed a GFP trap (Figure 2B). This showed very robust co-precipitation of WDR82 with ZC3H4-GFP confirming the two as interaction partners.

**Figure 2:**
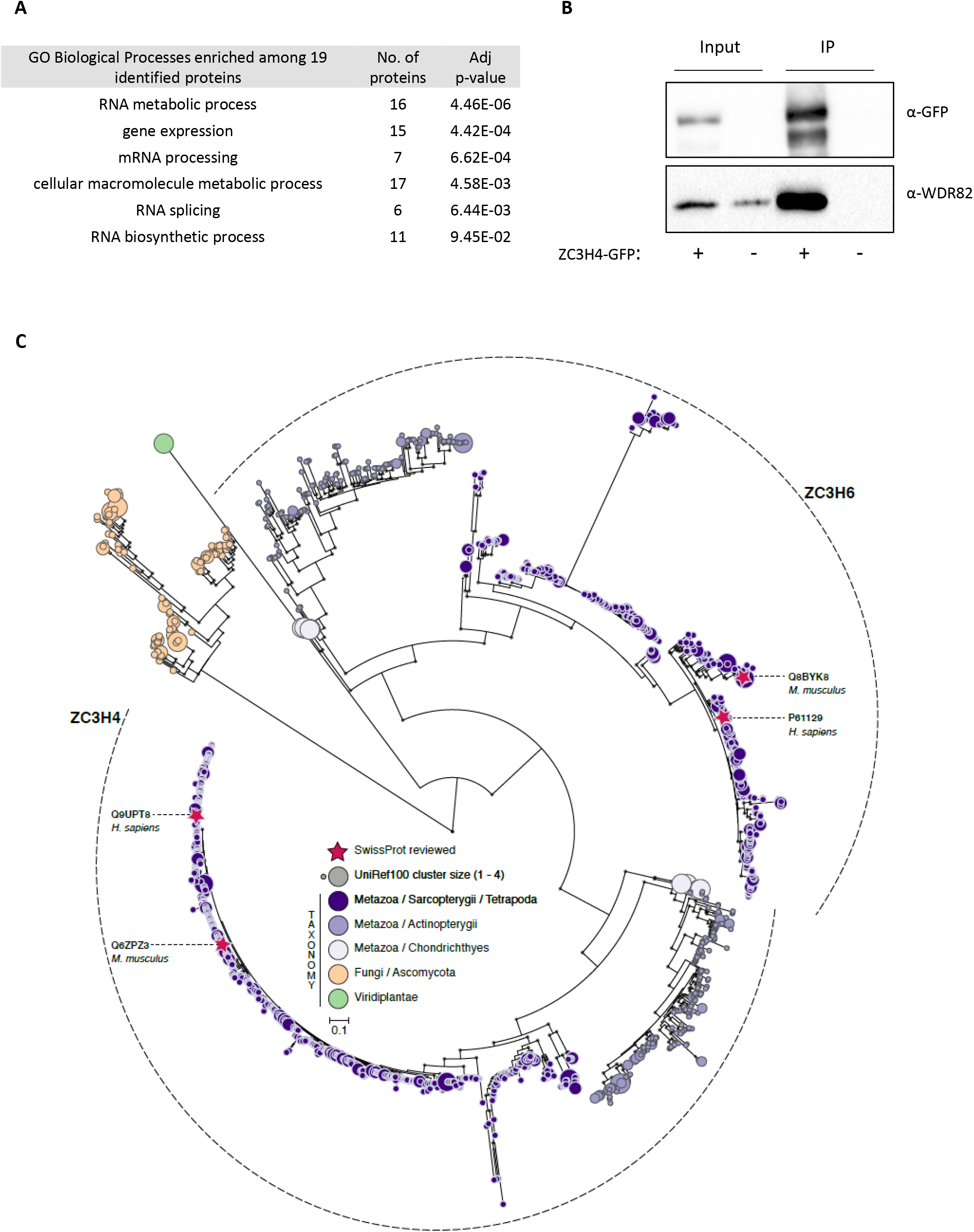
ZC3H4 binds the WDR82 termination factor and is metazoan-specific. a) Proteins that are co-regulated with ZC3H4 according to ProteomeHD using a score cut-off set to 0.98. Table shows GO term analysis of the potentially co-regulated factors. b) Co-immunoprecipitation of WDR82 using ZC3H4-GFP as bait. Blot shows input (5%) and immunoprecpitated material probed with antibodies to WDR82 or GFP. Cells untransfected with ZC3H4-GFP act as a negative control. c) Maximum-likelihood phylogenetic tree of zinc finger CCCH-domains (1513 sequences; 795 parsimony informative sites) inferred under the JTT+R8 model. Clades of ZC3H4-like and ZC3H6-like domains are delimited by dashed lines. CCCH-domains identified using the PANTHER hidden Markov model PTHR13119 against the UniProtKB protein database (non-redundant version: UniRef100; external node size represents protein cluster size). Branch support values ≥90% (based on 1000 ultrafast bootstraps) are indicated by grey circles. Red stars show SwissProt reviewed protein sequences; external nodes are color-coded according to their taxonomic lineage. Scale bar represents the number of estimated substitutions per site.

WDR82 is conserved between human and budding yeast. To determine the evolutionary background of ZC3H4, we conducted a phylogenetic analysis using a large sampling of homologous sequences (Figure 2C). These were identified using hidden Markov model (HMM)-based searches, corresponding to a zinc finger CCCH domain within ZC3H4, against the non-redundant protein database UniRef100. This identified 1513 UniRef100 sequences matching the HMM (Supplementary Excel file 2). ZC3H4 and ZC3H6-like sequences were recovered from this search, including SwissProt reviewed entries for *H. sapiens* and *M. musculus* for ZC3H4 (Q9UPT8 and Q6ZPZ3) and ZC3H6 (P61129 and Q8BYK8). Virtually all recovered sequences were from metazoan organisms—except for a group of fungal sequences from ascomycetes. A maximum-likelihood phylogeny was reconstructed using the identified zinc finger CCCH domain sequences. The resulting phylogenetic tree shows the dichotomy between the ZC3H4 and ZC3H6 domains, which are found in the same set of organisms. This indicates that they are paralogues and have likely diverged their function following gene duplication. The ancestral gene coding for ZC3H4/6 was likely lost from the non-vertebrates and subsequently underwent a duplication event leading to the ZC3H4- and ZC3H6-like paralogues in vertebrates.

### ZC3H4 restricts non-coding transcription events

To assess any function of ZC3H4 and/or ZC3H6 in RNA biogenesis we depleted either or both from HCT116 cells using RNA interference (RNAi) (Figure S3A), then deep sequenced nuclear transcripts. Comparison of these datasets shows that ZC3H4 loss has a more noticeable impact than ZC3H6 depletion (Figure S3B). Specifically, ZC3H6 depleted samples are more similar to control than those deriving from ZC3H4 loss and ZC3H4/ZC3H6 co-depletion resembles a knock-down of just ZC3H4. This was also evident from closer inspection of the data (Figure S3C), supporting the phylogenetic prediction of their separate functions. Accordingly, subsequent analyses focus on ZC3H4.

Due to its links with CPSF30 and WDR82, we anticipated that ZC3H4 might affect transcriptional termination. We first checked protein-coding genes and found a small number with longer read-through beyond the PAS when ZC3H4 is depleted (Figure 3A). However, broader analysis suggests that this is not widespread (Figure 3B and S3D). Visual inspection of the data instead revealed a strong effect of ZC3H4 depletion on PROMPTs. These antisense transcripts are short and rapidly degraded 3’→5’ by the exosome (Preker et al., 2008). Figure 3C shows an example PROMPT, upstream of *MYC*, which is undetectable in control siRNA treated cells, but abundant following ZC3H4 depletion. Loss of ZC3H4 also leads to a substantial extension of this transcript for more than 100 kilobases. This is made clearer by comparing the loss of ZC3H4 to AID-mediated depletion of the catalytic exosome (DIS3) (Davidson et al., 2019). This condition stabilises a shorter PROMPT RNA typical of early termination and rapid degradation. These effects are not isolated because meta-analysis reveals similar effects at many other PROMPTs (Figure 3D). This global comparison of ZC3H4 and DIS3 depletion datasets further illustrates the extended PROMPT transcription following depletion of the former.

**Figure 3:**
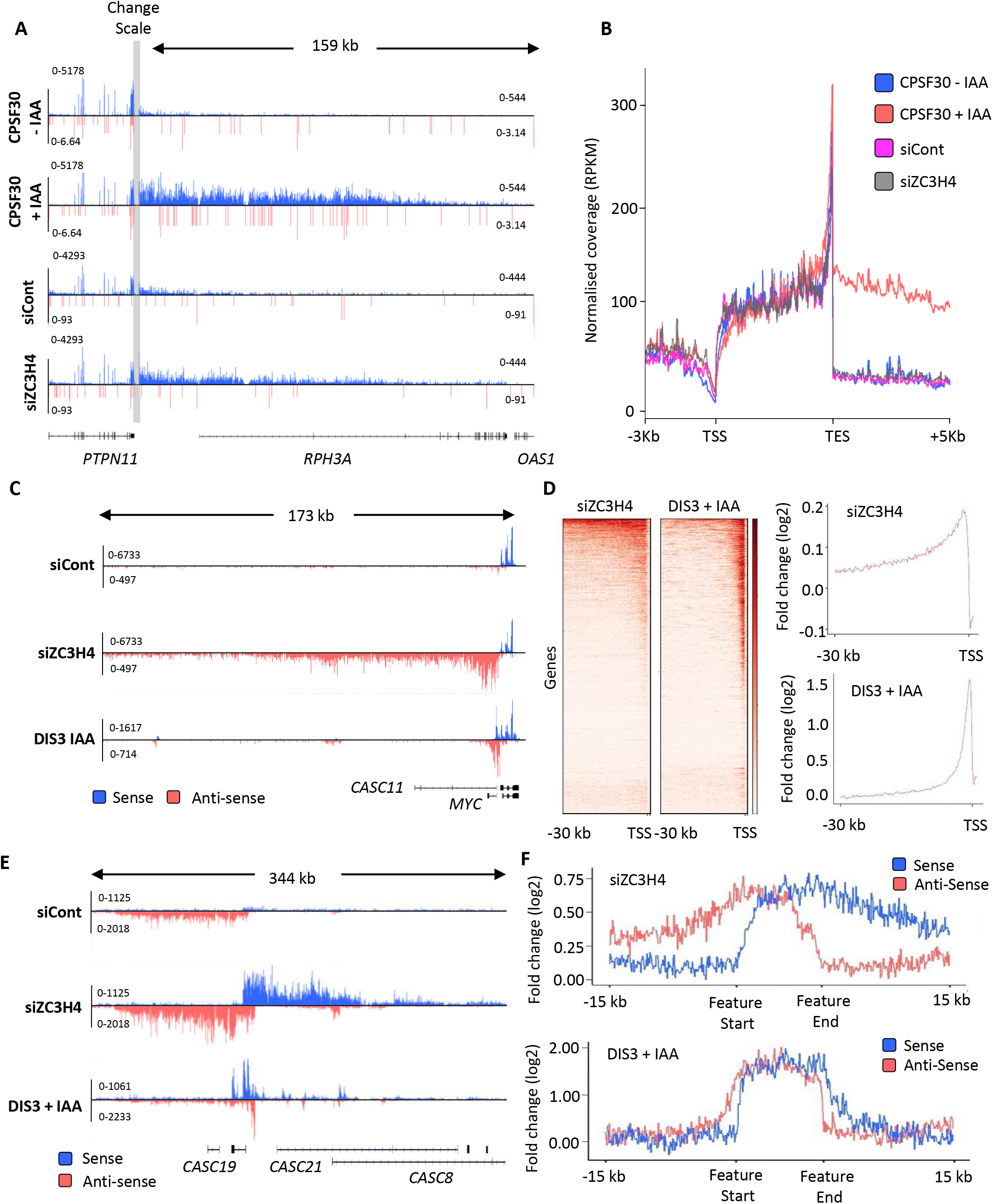
ZC3H4 depletion stabilises unproductive transcripts. a) IGV track of the transcription read-through defect at *PTPN11* following CPSF30 or ZC3H4 depletion. Blue and red tracks indicate sense/anti-sense transcripts respectively, grey bar indicates a change in y-axis scale so that comparatively weaker read through signals can be visualised next to the gene body (left scale for upstream of TES; right for downstream). Y-axis scale is RPKM. b) Metagene comparison of transcription upstream, across, and downstream of, protein-coding genes in nuclear RNA from *CPSF30-mAID* cells treated or not with auxin and from HCT116 cells transfected with control or ZC3H4 siRNAs. CPSF30 traces are from the same samples presented in Figure 1C. c) IGV track view of transcription at the *MYC* PROMPT in RNA-seq samples obtained from control or ZC3H4 siRNA treated HCT116 cells. We also show a track from HCT116 cells acutely depleted of DIS3-AID (Davidson et al., 2019) to highlight the normal extent of this transcript. Y-axis scale is RPKM. d) Log_2_ fold change of siZC3H4 vs siControl or DIS3 + vs −auxin for 6057 non-neighbouring, actively transcribed genes, plotted as heat maps. Line graphs are an XY depiction of heat map data. Log_2_ fold changes are smaller in siZC3H4 samples versus DIS3 depletion because this is an average of all genes in the heat map, a smaller fraction of which are affected by ZC3H4. e) IGV plot of a known SE upstream of *MYC.* Samples are shown from HCT116 cells treated with control or ZC3H4 siRNAs as well as *DIS3-AID* cells treated with auxin (the latter from (Davidson et al., 2019)) to show the normal extent of eRNAs over this region. Y-axis scale is RPKM. f) Log_2_ fold change of siZC3H4 vs siControl or DIS3 + vs −auxin for 111 SEs. The bed file detailing super-enhancer coordinates in HCT116 cells was taken from dbSUPER.org.

The finding that PROMPTs are affected by ZC3H4 suggested a role in the transcription/metabolism of antisense/non-coding RNAs. We therefore extended our search for potential ZC3H4 regulated transcription to enhancer regions since they also produce short RNAs that are degraded by the exosome (Andersson et al., 2014). eRNAs can be found in isolation and in clusters called super-enhancers (SEs) (Pott and Lieb, 2015). SEs are thought to be important for controlling key developmental genes with strong relevance to disease (Hnisz et al., 2013). ZC3H4 depletion has a clear effect over SE regions exemplified by the *MYC* SE in where upregulation and extension of eRNAs is obvious (Figure 3E). Again, acute depletion of DIS3 illustrates the normally restricted range of individual eRNAs within the cluster. This effect is general for other SEs as demonstrated by the metaplots in Figure 3F. We also checked the effect of CPSF30 depletion on example PROMPT and SE transcription, which are very modest (Figure S3E). Overall, these data strongly suggest that ZC3H4 is important for regulating transcription across many PROMPTs and SEs.

### Comparison of ZC3H4 and Integrator effects

ZC3H4 has some functions in common with the Integrator complex, being metazoan-specific and relevant to the metabolism of unstable non-coding RNA (Lai et al., 2015; Mendoza-Figueroa et al., 2020; Nojima et al., 2018). We previously published chromatin RNA-seq derived from cells RNAi depleted of the Integrator backbone component INTS1 (Davidson et al., 2020). Metagene analysis of this data at protein-coding genes shows a mild effect of Integrator depletion over PROMPT regions (Figure 4A). It also reveals an accumulation of promoter-proximal RNAs in the coding direction consistent, which fits well with a recent report on its function as an attenuator of protein-coding transcription (Lykke-Andersen et al., 2020).

**Figure 4:**
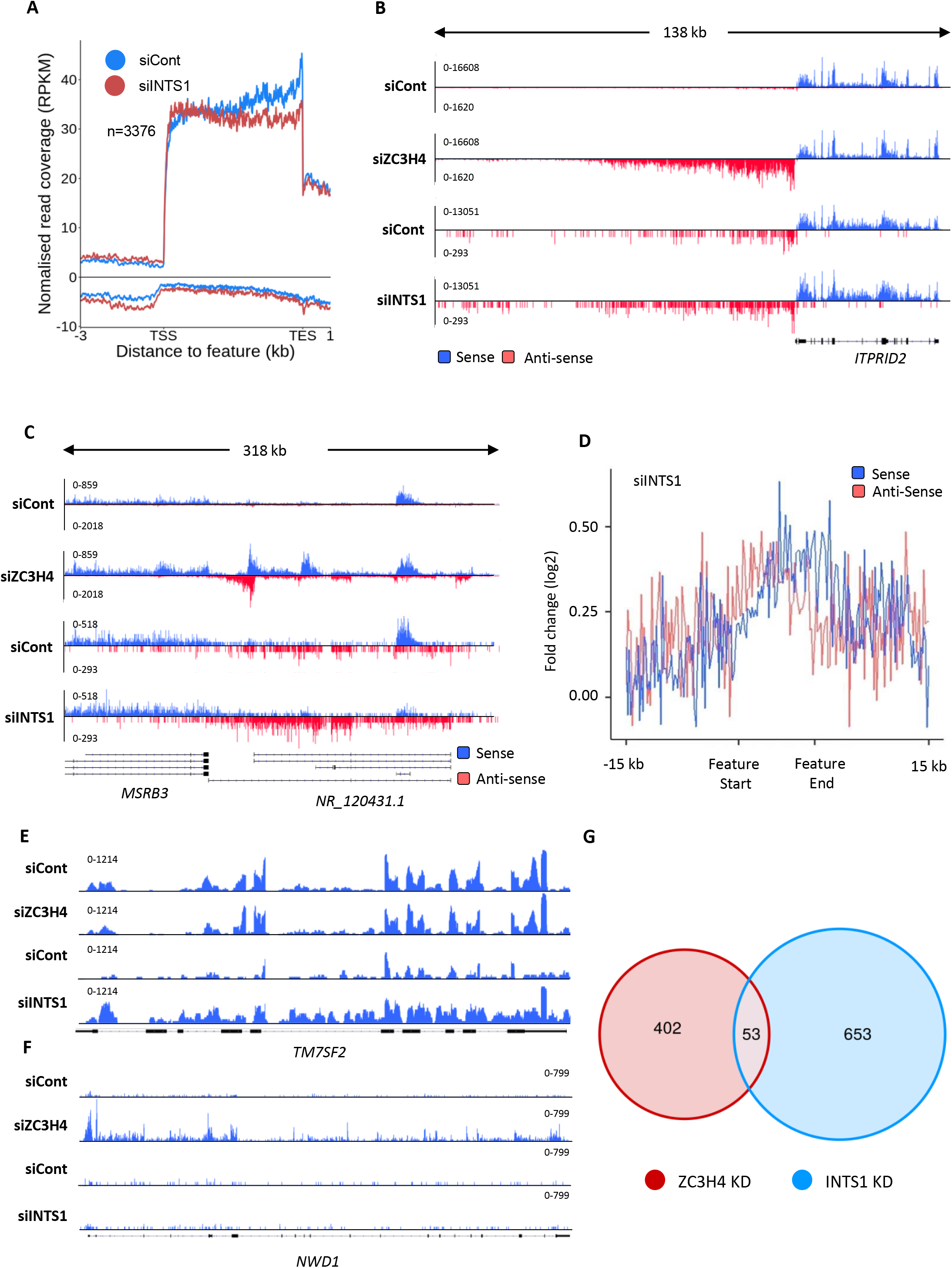
Comparison of ZC3H4 and Integrator effects. a) Metagene analysis of chromatin-associated RNA-seq performed on cells treated with control or INTS1-specific siRNA. The plot shows signals upstream, across and downstream of protein-coding genes. Signals below the line are antisense transcription. Y-axis scale is RPKM. b) Comparison of chromatin-associated RNA-seq in control and INTS1 siRNA treated samples with nuclear RNA-seq derived from control or ZC3H4 siRNA treatment. The *ITPRID2* PROMPT is displayed and y-axes are RPKM (note the different scales between ZC3H4 and INTS1 samples). c) Comparison of chromatin-associated RNA-seq in control and INTS1 siRNA treated samples with nuclear RNA-seq derived from control or ZC3H4 siRNA treatment. The *MSRB3* SE is displayed and y-axes are RPKM (note the different scales between INTS1 and ZC3H4 samples). d) Metaplot of RNA-seq profile over super-enhancers following INTS1 depletion. Log_2_ fold over 111 super-enhancer as line graphs. The bed file detailing super-enhancer coordinates in HCT116 cells was taken from dbSUPER.org. e/f) IGV traces of *NWD1* and *TM7SF2* genes derived from chromatin-associated RNA-seq in control and INTS1 siRNA treated samples and nuclear RNA-seq from control or ZC3H4 siRNA treatment. *NWD* transcripts are affected by ZC3H4 but not INTS1 whereas the opposite is true for *TM7SF2* RNAs. Y-axes scales are RPKM. g) Venn diagram showing the number of mRNAs upregulated ≥ 2-fold, padj ≤0.05 following ZC3H4 depletion versus INTS1 loss and the overlap between the two sets. Genes that showed increased expression due to transcription read-through from an upstream gene were also discarded by assessing coverage over a 1 kb region preceding the TSS, relative to untreated cells.

Where ZC3H4 effects are evident over PROMPT regions, they are generally more substantial than those seen after Integrator loss, exemplified by the *ITPRID2* PROMPT in Figure 4B. Likewise, at SEs, ZC3H4 depletion generally results in a greater stabilisation and elongation of eRNA, compared to INTS1 knock-down, exemplified at the *MSRB3* SE (Figure 4C). Meta-analysis confirms this difference (compare Figures 4D and 3F). We note that INTS1 data are on chromatin-associated RNA whereas ZC3H4 images are obtained from nuclear RNA. However, as chromatin-associated RNA is more enriched in nascent transcripts this would be expected to capture more extended non-coding transcription and not less. Therefore, it is unlikely that the lower effects of Integrator are accounted for by this technical difference.

Another important role of Integrator is in the promoter-proximal termination of protein-coding gene transcription (Elrod et al., 2019; Lykke-Andersen et al., 2020; Tatomer et al., 2019). Increased full-length transcription of a subset of genes is seen in its absence reflecting escape from this early termination. *TM7SF2* is an example of a gene where this is seen following INTS1 depletion (Figure 4E). We found similar effects at other genes (e.g. *NWD1*) when ZC3H4 is depleted (Figure 4F). Interestingly, there was little overlap between the protein-coding transcripts upregulated following INTS1 or ZC3H4 loss (Figure 4E, Supplementary Excel file 3). This reflects the interesting possibility that they influence the transcription of distinct gene sets.

### Rapid ZC3H4 depletion recapitulates RNAi effects

ZC3H4 RNAi suggests its widespread involvement in non-coding RNA synthesis. However, RNAi depletion was performed using a 72 hr protocol and might result in indirect or compensation effects. To assess whether these effects are a more direct consequence of ZC3H4 loss, we engineered HCT116 cells for its rapid and inducible depletion. CRISPR/Cas9 was used to homozygously tag *ZC3H4* with an *E.coli* derived DHFR degron preceded by 3xHA epitopes (Figure 5A; (Sheridan and Bentley, 2016)). In this system, cells are maintained in trimethoprim (TMP) to stabilise the degron, removal of which causes protein depletion. Western blotting demonstrates homozygous tagging of *ZC3H4* and that ZC3H4-DHFR is depleted following TMP removal (Figure 5B). Depletion was complete after overnight growth without TMP but substantial protein loss was already observed after 4 hrs allowing us to assess the consequences of more rapid ZC3H4 depletion.

**Figure 5:**
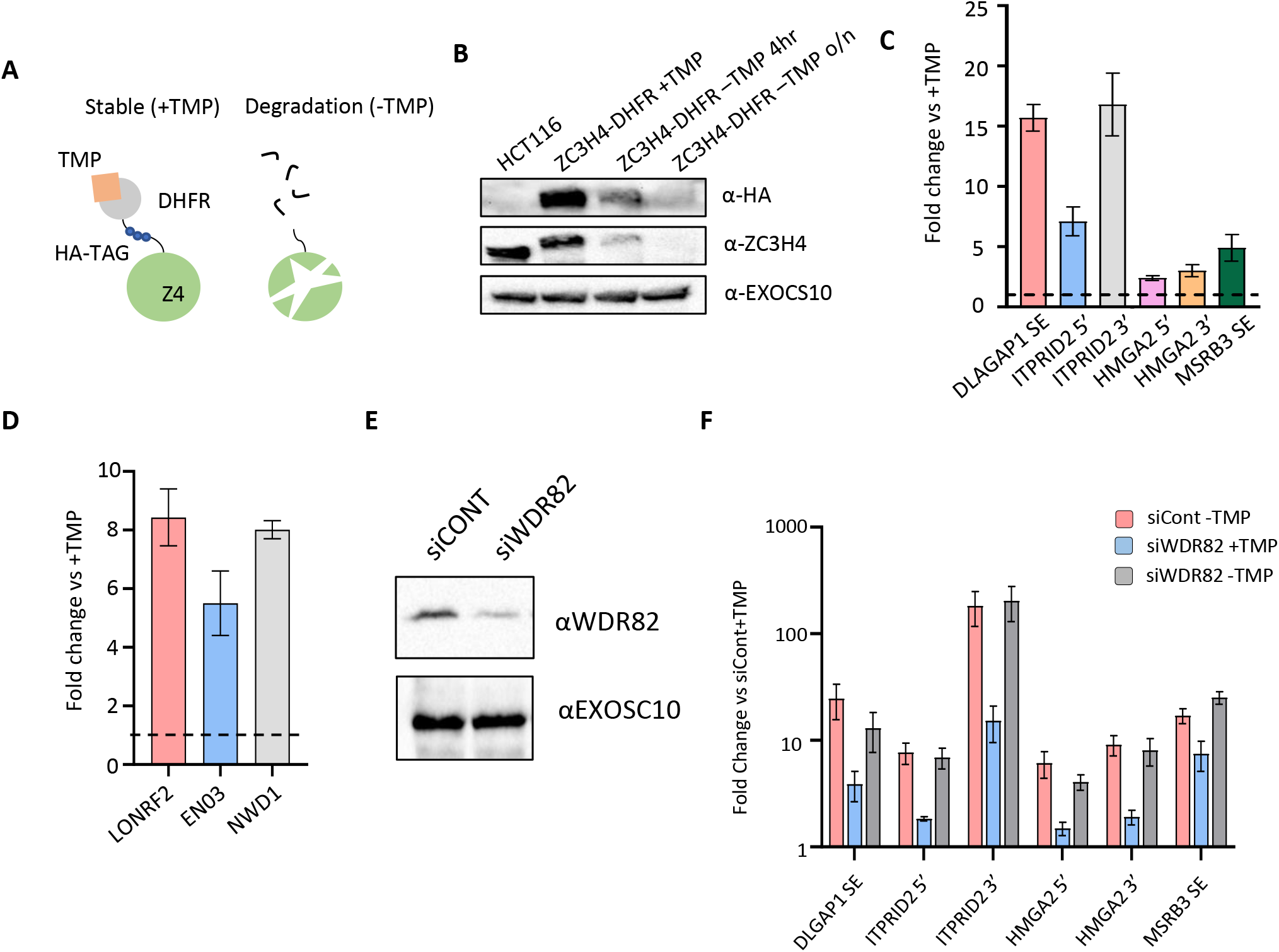
Transcriptional dysregulation following acute ZC3H4 loss. a) Schematic detailing how the DHFR degron works. *E. Coli* dihydrofolate reductase (DHFR) is fused to the C-terminus of ZC3H4, which is stabilised by 2,2,6,6-Tetramethylpiperidine (TMP). When TMP is removed ZC3H4-DHFR is degraded. b) Western blot of HCT116 parental and HCT116 ZC3H4-DHFR cells +/− TMP, TMP was withdrawn for 4 hours or overnight, EXOCS10 is used as a loading control, αHA recognises a HA peptide prior to the DHFR tag, while αZC3H4 recognises native protein sequence. c) qRT-PCR analysis of PROMPT and SE transcripts in *ZC3H4-DHFR* cells grown with or without (4hr) TMP. Graph shows fold change versus +TMP (showed by dotted line) following normalisation to spliced actin. N=4. Error bars are standard error of the mean (SEM). d) qRT-PCR analysis of spliced LONRF2, ENO3 and NWD1 mRNAs in *ZC3H4-DHFR* cells grown with or without (4hr) TMP. Graph shows fold change versus +TMP (showed by dotted line) following normalisation to spliced actin. N=3. Error bars are SEM. e) Western blot of extracts derived from HCT116 cells transfected with control or WDR82-specific siRNAs. The blot shows WDR82 and, as a loading control, EXOSC10. f) qRT-PCR of PROMPT and SE transcripts in *ZC3H4-DHFR* cells transfected with control or WDR82 siRNAs before withdrawal, or not, of TMP (14hr). Graph shows fold change by comparison with control siRNA transfected *ZC3H4-DHFR* cells maintained in TMP following normalisation to spliced actin transcripts. N=3. Error bars are SEM.

*ZC3H4-DHFR* cells were then grown in the presence or absence of TMP (4 hrs) before isolation of total RNA. This was analysed by qRT-PCR to assess the levels of extended PROMPT (*HMGA2, ITPRID2*) and SE (*MSRB3, DLGAP1*) RNAs (Figure 5C). All were increased following ZC3H4 loss suggesting that the effects that we observed by RNA are not due to compensatory pathways. We also analysed three protein-coding transcripts (ENO3, NWD1 and LONRF2) that showed upregulation following ZC3H4 RNAi (Figure 5D). All showed 6-8 fold upregulation after 4hr ZC3H4 depletion suggesting a role in controlling the output from these genes.

Figure 2B demonstrated that ZC3H4 interacts with WDR82. To test whether WDR82 is involved in the transcription of ZC3H4 targets we depleted it from *ZC3H4-DHFR* cells (Figure 5E). Since WDR82 is present in multiple protein-complexes, we wished to assess whether it is involved in alternative/additional termination pathways at ZC3H4-affected loci. As such, we co-depleted WDR82 and ZC3H4, reasoning that a synergistic effect would indicate WDR82-dependent processes that do not involve ZC3H4 (Figure 5F). WDR82 depletion caused upregulation of all ZC3H4 targets consistent with it being part of the same pathway. There was no synergistic effect of their co-depletion suggesting that WDR82 and ZC3H4 share a common pathway at these loci. Finally, we depleted either PNUTS or SETD1A/B since they form separate complexes with WDR82. Loss of the former but not the latter affected transcript levels at ZC3H4-affected loci (Figure S4A-C).

### ZC3H4 occupies a broad region at a subset of promoters

We have demonstrated that depletion of ZC3H4 causes widespread defects in non-coding transcription and at a subset of protein-coding genes. As these effects are seen following rapid ZC3H4 depletion, we anticipated them to be direct. Given the acute instability of PROMPTs and eRNAs, we hypothesised that recruitment of ZC3H4 to these loci would underpin its effects. The proximity of ZC3H4 to transcription is supported by its capture in our mTurbo experiment and by the presence of CCCH zinc finger domains that are predicted to bind nucleic acid (Figure S2A). To assay its genomic occupancy, we performed ZC3H4 chromatin immunoprecipitation and sequencing (ChIP-seq) alongside that of Pol II.

ZC3H4 occupies genes with binding broadly resembling that of Pol II and showing the greatest enrichment over promoter regions (Figure 6A). However, many genes that are occupied by Pol II do not recruit ZC3H4 (Figure 6B). This might result from poorer affinity of the ZC3H4 antibody or that its recruitment to chromatin is bridged, since ZC3H4 also directly crosslinks to RNA in cells (Figure S5A). However, its differential gene occupancy is consistent with the RNA-seq data showing ZC3H4 depletion to impact only certain transcripts. Furthermore, ZC3H4 occupies a broader promoter region than Pol II suggesting that its function is not restricted to the precise transcriptional start site. The width of this peak often corresponds to the normal extent of PROMPT and eRNA transcription (1-2kb), which is elongated in its absence. *RPL13* is shown as an example of recruitment of ZC3H4 upstream of the promoter, where its loss causes stabilisation and extension of the antisense transcript (Figure 6C). ZC3H4 is also strongly recruited to SEs consistent with the RNA effects observed on them following its loss (Figure 6D). This is exemplified by the *MSRB3* region and generalised by metaplots in figure 6E. Although our analyses of eRNA and PROMPTs was guided by our RNA-seq findings, an unbiased search for peaks of ZC3H4 and Pol II signal confirmed proportionally greater ZC3H4 occupancy at distal intergenic regions (encompassing SEs) (Figure 6F).

**Figure 6:**
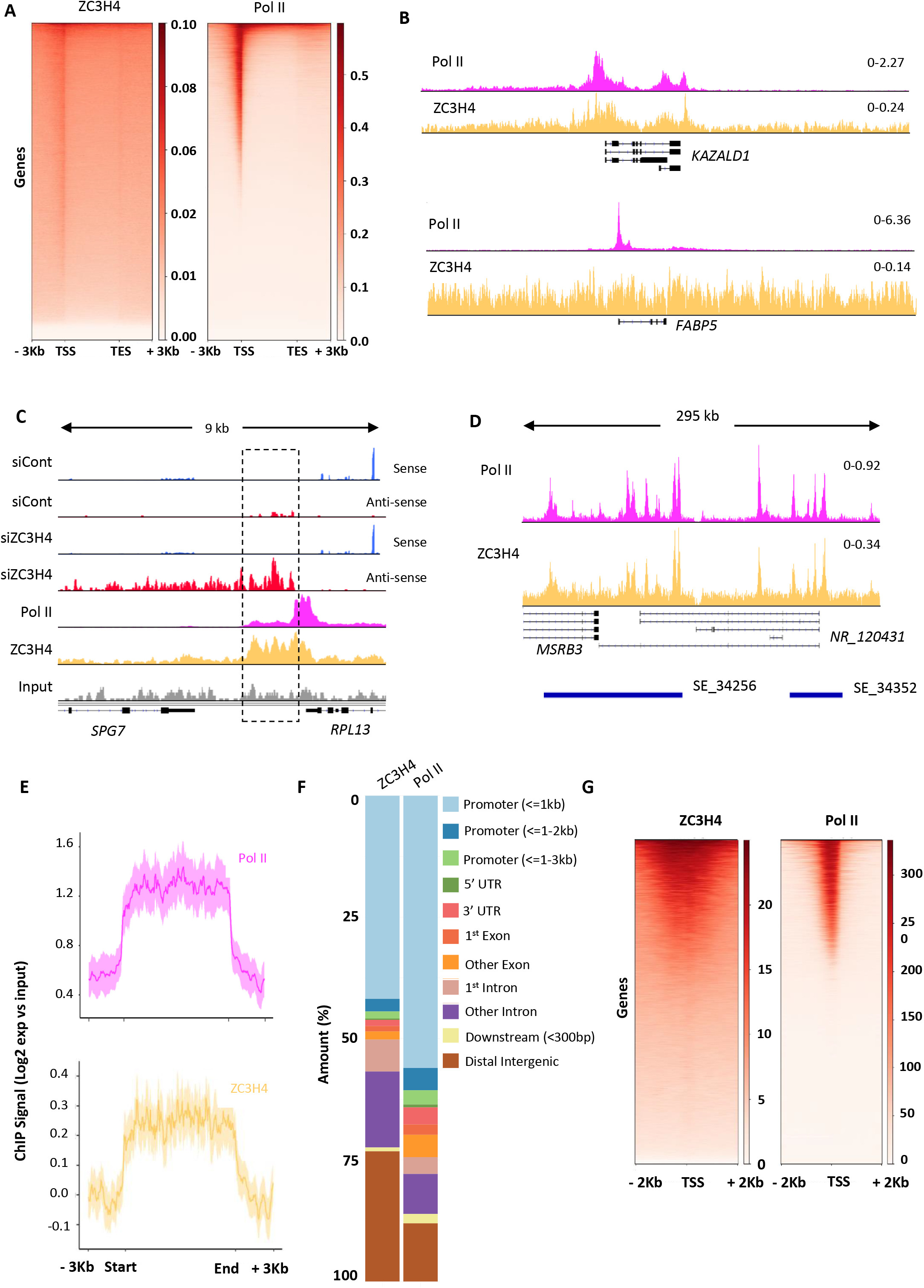
ZC3H4 ChIPs occupies regions of observable transcriptional dysregulation. a) ZC3H4 ChIP profile over protein-coding genes is similar to Pol II. Heat map representation of ZC3H4 and Pol II ChIP-seq occupancy over the gene body +/− 3 kb. b) ZC3H4 occupies fewer promoters than Pol II. IGV track view of ZC3H4 and Pol II occupancy over *KAZALD1* and *FABP5* genes, Pol II is present at both genes, while ZC3H4 is only present at *KAZALD1*. Scale is counts per million (CPM). c) RNA-seq (HCT116 cells treated with control or ZC3H4 siRNA) and ChIP-seq (Pol II, ZC3H4 and input) profiles at *RPL13.* ZC3H4 occupancy is focused more on the PROMPT transcript region (hatched box) than the TSS where, in contrast, Pol II signal is maximal. RNA-seq scale is RPKM and ChIP-seq is CPM. Dotted box demarks approximate extent of PROMPT transcription referred to in the text. d) ZC3H4 ChIP occupancy mirrors Pol II at super-enhancers. IGV track view of ZC3H4 and Pol II occupancy over the SE at the *MSRB3* locus. HCT116 super-enhancer gene track is from dbSUPER and depicted as blue bars. e) Log_2_ fold change of ZC3H4 and Pol II vs input at SEs shown as a line graph. Halo denotes 95% confidence level. f) ChIPseeker analysis of peak distribution of ZC3H4 and Pol II. Occupancy regions are colour-coded and the number of ChIP peaks expressed as a proportion of 100%. g) Heat map showing Pol II and ZC3H4 ChIP occupancy in HEPG2 cells obtained via the ENCODE consortium. Occupancy +/− 2 kb of the TSS is shown.

Overall, the HCT116 ChIP-seq demonstrates direct recruitment of ZC3H4 to potential targets. One caveat is the low ChIP efficiency of the ZC3H4 antibody; however, a ZC3H4 ChIP-seq experiment was recently made available by the ENCODE consortium (Partridge et al., 2020). This used a flag-tagged construct and was performed in HEPG2 cells allowing a comparison of our data to that obtained with a high-affinity antibody and in different cells. The data are consistent with our findings for endogenous ZC3H4 in HCT116 cells. Flag-ZC3H4 occupies a subset of Pol II-bound regions and shows broader distribution than Pol II around promoters (Figure 6G). Although HEPG2 cells express fewer SEs than HCT116 cells, the transcribed *DLGAP1* example confirms its occupancy of these regions in both cell types (Figure S5B). In contrast, the *MYC* SE is only expressed in HCT116 cells and is not occupied by ZC3H4 in HEPG2 cells. In further agreement with our data, bioinformatics assignment of flag-ZC3H4 binding sites yielded “promoter and enhancer-like” as the most enriched terms (Partridge et al., 2020).

### Engineered recruitment of ZC3H4 suppresses transcription

The consequences of ZC3H4 recruitment to targets are predicted to be their early termination and subsequent degradation by the exosome, based on the well-established fate of PROMPTs and eRNAs. To test whether ZC3H4 recruitment can promote these effects, we established a tethered function assay. ZC3H4 was tagged with bacteriophage MS2 coat protein to engineer its recruitment to a reporter containing MS2 hairpin binding sites (MS2hp-IRES-GFP; Figure 7A). To control for non-specific effects, we employed MS2-GFP and to assess any effect of ZC3H4 over-expression we used ZC3H4-GFP. HCT116 cells were transfected with either of these three constructs together with MS2hp-IRES-GFP and reporter expression assayed by qRT-PCR. Compared to the two controls, tethered ZC3H4-MS2 dramatically reduced reporter RNA expression (Figure 7B). Importantly, ZC3H4-MS2 expression has no clear effect on the same reporter lacking MS2 hairpins (Figure 7C). This directly demonstrates that ZC3H4 recruitment is sufficient to negatively regulate RNA expression, mirroring the upregulation of its endogenous targets seen when it is depleted.

**Figure 7:**
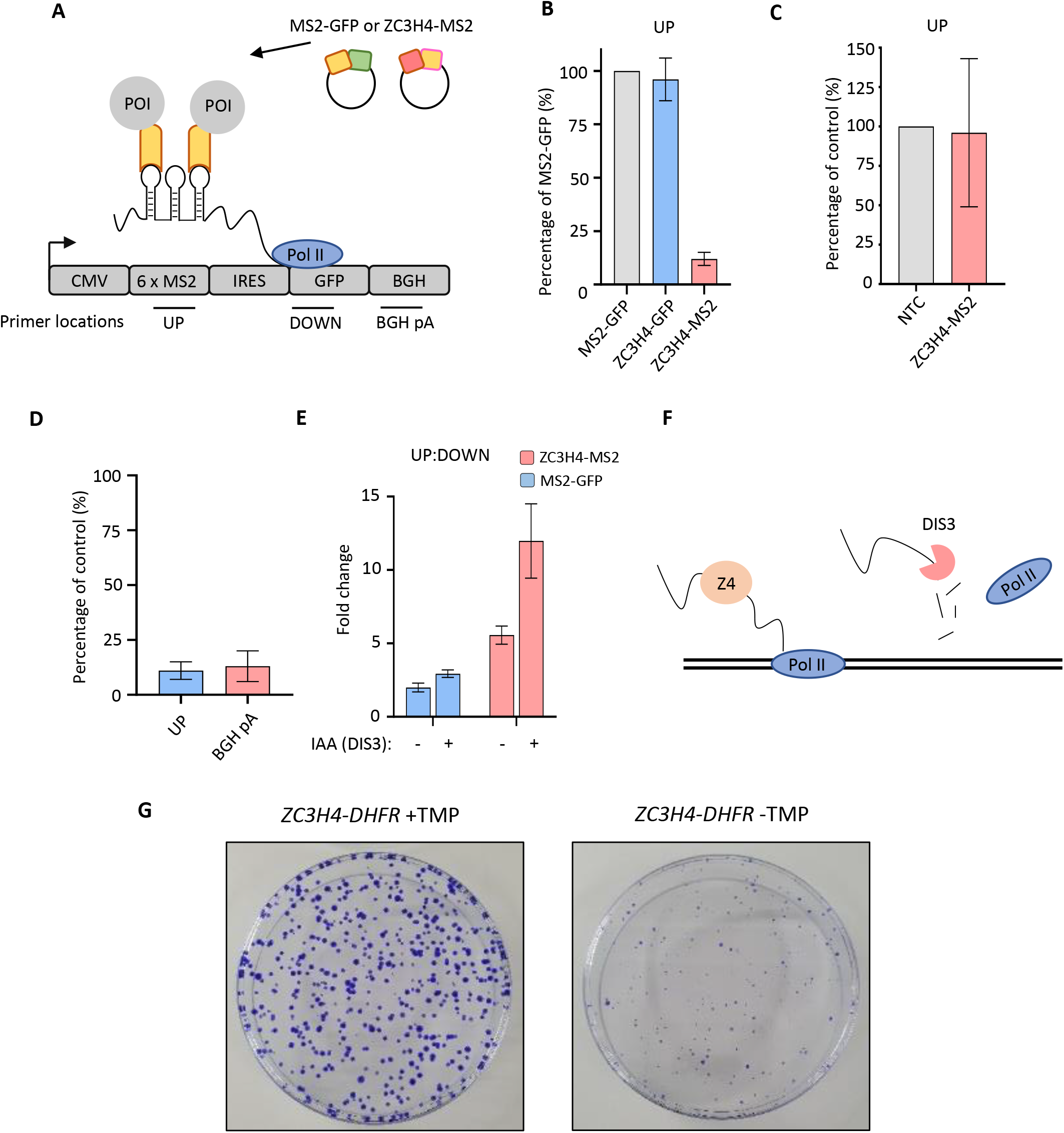
Directed recruitment of ZC3H4 recapitulates its effects on endogenous targets. a) Schematic of the MS2 system. A reporter plasmid (MS2hp-IRES-GFP) expressing a GFP transcript with 6 x MS2 hairpins upstream of an IRES and GFP gene. ZC3H4-MS2 or MS2-GFP can be specifically tethered to the MS2 hairpins to assess consequent effects on transcription/RNA output. Positions of primer pairs used in qRT-PCR experiments elsewhere in the figure are indicated by labelled horizontal lines under reporter. POI is protein of interest. b) qRT-PCR analysis of total RNA isolated from MS2hp-IRES-GFP transfected cells co-transfected with either MS2-GFP, ZC3H4-GFP or ZC3H4-MS2. The level of reporter RNA is plotted (“UP” amplicon) as a percentage of that obtained in the MS2-GFP sample following normalisation to spliced actin. N=3. Error bars are SEM. c) qRT-PCR of HCT116 cells transfected with IRES-GFP and either a control beta-globin plasmid (NTC) or ZC3H4-MS2. Graph shows percentage of GFP RNA versus control following normalisation to spliced actin transcripts. N=3. Error bars show SEM. d) qRT-PCR analysis of chromatin-associated RNA isolated from MS2hp-IRES-GFP transfected cells co-transfected with either MS2-GFP or ZC3H4-MS2. The level of reporter RNA upstream of the MS2 hairpins (UP) and transcripts yet to be cleaved at the BGH poly(A) site (BGH pA) are plotted as a percentage of that obtained in the MS2-GFP sample following normalisation to spliced actin. N=3. Error bars are SEM. e) qRT-PCR analysis of total RNA isolated from MS2hp-IRES-GFP transfected *DIS3-AID* cells co-transfected with either MS2-GFP or ZC3H4-MS2 – simultaneously treated or not with auxin (14hr in total). The graph shows the ratio of RNA species recovered upstream (UP) versus downstream (DOWN) of the MS2 hairpins. N=3. Error bars are SEM. f) Schematic detailing an interplay between ZC3H4 and DIS3 that sees transcription stop and nascent RNA degraded g) Colony formation assay of *ZC3H4-DHFR* cells grown in the presence or absence of TMP. Cells were grown for 10 days before crystal violet staining.

PROMPTs and eRNAs are degraded on chromatin and we wanted to test whether ZC3H4-MS2 exerted its effect on these nascent RNAs. Chromatin-associated RNA was isolated from cells transfected with MS2hp-IRES-GFP and either ZC3H4-MS2 or MS2-GFP. We also included an additional primer set to detect RNA uncleaved at the bovine growth hormone (BGH) poly(A) site. Because poly(A) site cleavage is co-transcriptional, this primer set should robustly detect Pol II-associated transcripts. This amplicon and that over the MS2 region were reduced in this chromatin fraction, strongly suggesting that tethered ZC3H4 acts on nascent RNA (Figure 7D).

The exosome targets released PROMPT and eRNA transcripts, which could be influenced by ZC3H4. The results we present for endogenous loci are consistent with this since PROMPTs and eRNAs are upregulated and elongated when ZC3H4 is depleted. To test whether recruited ZC3H4 leads to exosome decay, we transfected MS2hp-IRES-GFP, together with either MS2-GFP or ZC3H4-MS2, into *DIS3-AID* cells that were then treated or not with auxin to eliminate the catalytic exosome. RNA upstream and downstream of the MS2hp region was detected by qRT-PCR and their ratio plotted (Figure 7E). Enhanced levels of upstream versus downstream amplicon was associated with transfection of ZC3H4-MS2 and depletion of DIS3. This is consistent with the hypothesis that recruited ZC3H4 promotes the release of RNA that is a DIS3 substrate (Figure 7F).

Finally, having demonstrated the importance of ZC3H4 in transcription, we were interested to determine its overall relevance to cell growth. This is made simpler by the *ZC3H4-DHFR* cell line, which allows permanent depletion of ZC3H4 by growth without TMP. Accordingly, we performed colony formation assays on these cells grown in the presence or absence of TMP (Figure 7G). Loss of ZC3H4 was associated with smaller colonies clearly demonstrating the importance of ZC3H4 for growth/proliferation.

## DISCUSSION

We have discovered that ZC3H4 controls unproductive transcription with non-coding transcripts a prominent target. This conclusion is based on its recruitment to loci that give rise to transcripts that are stabilised and elongated when it is depleted. Moreover, engineered recruitment of ZC3H4 to a heterologous reporter is sufficient to promote its degradation by the exosome. We propose that ZC3H4 recruitment drives some of the early transcriptional termination that is characteristic of many non-coding RNAs, particularly PROMPT and eRNA transcripts. The function of ZC3H4 in restraining their transcription may at least partly explain why PROMPT and eRNA transcripts accumulate as short species when the exosome is depleted.

Another non-coding RNA termination mechanism involves the Integrator, which terminates PROMPTs and eRNAs (Lai et al., 2015; Nojima et al., 2018). We compared the effects of Integrator and ZC3H4 at both transcript classes. Where ZC3H4 effects are apparent, they show greater stability and longer extensions in its absence versus when Integrator is depleted. One explanation is that Integrator and ZC3H4 act at different stages in non-coding RNA biogenesis. Although we cannot rule out that they act redundantly, ZC3H4 has not been found in complex with Integrator as far as we know. Moreover, the protein-coding transcripts upregulated by ZC3H4 and Integrator loss show little overlap suggesting that they are controlled by independent mechanisms. This raises the exciting possibility that multiple factors may control promoter-proximal termination at distinct gene sets.

While ZC3H4 clearly affects some protein-coding transcription, its biggest impact is at unstable non-coding RNAs; however, again this is a subset of the total. This could be because RNA-seq was performed after ZC3H4 RNAi rather than following acute depletion. However, this is unlikely because ZC3H4 is recruited to the loci where transcription is affected in its absence. Precisely how ZC3H4 is recruited to its targets requires elucidation. The TurboID experiment demonstrates its proximity to Pol II and IP experiments show it to bind WDR82, which also affects ZC3H4-dependent transcripts. WDR82 interacts directly with Pol II that is phosphorylated on Serine 5 of its C-terminal domain (Lee and Skalnik, 2008), and could play a role in recruiting ZC3H4 to promoter-proximal regions. We also show that ZC3H4 UV cross-links to RNA in cells, which may also underpin its observed selective recruitment to certain loci (Figure S5A). Lastly, ZC3H4 has been bioinformatically identified as a direct DNA binder by ENCODE (Partridge et al., 2020). Future experiments will be directed toward dissecting the nucleic acid binding capabilities of ZC3H4 and their relevance to its recruitment and activities.

Although very little is published on ZC3H4 itself, it has been proposed as an equivalent to *Drosophila* Suppressor of Sable (Su(s)), which negatively regulates transcription via promoter-proximal termination (Brewer-Jensen et al., 2016; Kuan et al., 2004). ZC3H4 and Su(s) share little sequence homology, evidenced by the absence of Su(s) in our phylogenetic analysis. However, they have similar structural makeup with zinc fingers flanked by largely disordered regions. Like ZC3H4, Su(s) depletion stabilises RNAs and causes their elongation.

Although a global analysis of Su(s) depletion has not been carried out, example substrates are degraded by the exosome. Su(s) binds RNA and DNA, which may enable a variety of targeting mechanisms. Interestingly, IP and mass spectrometry indicates that WDR82 may be the only interacting partner of Su(s) (Brewer-Jensen et al., 2016). There is no known catalytic activity for ZC3H4 or Su(s). However, both are related to CPSF30, which shows endonuclease activity in *Drosophila* and *Arabidopsis* (Addepalli and Hunt, 2007; Bai and Tolias, 1996). It remains to be seen whether ZC3H4 possesses any catalytic activity or mediates its effects through interaction partners.

The strong ZC3H4 occupancy of SEs, together with the disruption of their transcription in its absence, is also interesting. Other notable SE-associated factors (e.g. BRD4 and MED1) are much more generally implicated in Pol II transcription than ZC3H4 (Sabari et al., 2018). Moreover, they are transcriptional activators whereas ZC3H4 appears to suppress transcription (or, at least, its RNA output). Many SE-bound factors are found to have phase separation properties explaining why large clusters of factors accumulate at these regions (Cho et al., 2018). While we do not know whether ZC3H4 can phase separate, it contains large regions of intrinsic disorder, which can promote this property. In general, ZC3H4 may offer a new way to study enhancer clusters, particularly the importance of restricting transcription across these regions.

In conclusion, we have uncovered ZC3H4 as a factor with a function in restricting transcription. Its most notable effects are at non-coding loci where transcriptional termination mechanisms are less understood than at protein-coding genes. The further dissection of ZC3H4 and its targeting should reveal additional important insights into how this unstable portion of the transcriptome is controlled. By uncovering non-overlapping effects of Integrator and ZC3H4 at protein-coding genes our data indicate the possibility that multiple factors may control higher eukaryotic gene output via premature transcriptional termination.

## ACKNOWLEDGEMENTS

We are grateful to the other members of the lab for critical comment. This work was supported by a Wellcome Trust Investigator Award (WT107791/Z/15/Z) and a Lister Institute Research Fellowship held by S.W. We are grateful to The University of Exeter Sequencing Service where all sequencing was performed who are supported by a Medical Research Council Clinical Infrastructure Award (MR/ M008924/1), the Wellcome Trust Institutional Strategic Support Fund (WT097835MF), a Wellcome Trust Multi User Equipment Award (WT101650MA), and a Biotechnology and Biological Sciences Research Council Longer and Larger (LoLa) Award (BB/ K003240/1).

## AUTHOR CONTRIBUTIONS

C.E performed most experiments and bioinformatics, prepared the figures and helped draft the manuscript. L.D performed some bioinformatics analysis as did P.S. A.M did the phylogenetic analysis. S.W performed the ZC3H4-DHFR depletion experiments, made and analysed the PNUTS-AID cell line, supervised the project and wrote the manuscript with input from all authors.

## EXPERIMENTAL PROCEDURES

### Cell culture

HCT116 parental cells and engineered cell lines were cultured in Dulbecco modified eagle medium, supplemented with 10% foetal calf serum and penicillin streptomycin (Gibco). For RNAi, 6 or 24-well dishes were transfected with siRNA using Lipofectamine RNAiMax (Life Technologies) following the manufacturers’ guidelines. The transfection was repeated 24 hours later and, 48 hours after the first transfection, RNA was isolated. For MS2 assays, cells were seeded in 24-well dishes overnight, then transfected with 50ng MS2hp-IRES-GFP and 100ng of MS2-GFP, ZC3H4-MS2 or ZC3H4-GFP using JetPRIME® (PolyPlus) for 24 hours. To deplete DIS3-AID, auxin was used at a final concentration of 500uM. To deplete ZC3H4-DHFR, cells were washed twice in PBS and grown in media with or without TMP (30mM).

### Cell line generation and cloning

*CPSF30-mAID* and *CPSF30-mAID*:*RPB1-mTurbo* cells lines were generated using CRISPR/Cas9-mediated homology-directed repair (HDR). CPSF30 and RPB1 homology arms and gRNA sequences are detailed in Supplementary excel file 4. The mTurbo insert derives from 3xHA-mTurbo-NLS_pCDNA3 (#107172, Addgene). For ZC3H4 degron cells, 3xHA-DHFR was amplified from existing CPSF73-HA-DHFR constructs (published in (Eaton et al., 2018)) using non-homologous end-joining (NHEJ) as described in (Manna et al., 2019). PNUTS-AID cells were constructed using the protocol described in (Davidson et al., 2019). In general, 6cm dishes of cells were transfected with 1ug of guide RNA expressing px300 plasmid (#42230, Addgene) and 1ug of each HDR template/NHEJ PCR product. Three days later cells were seeded, as appropriate, into hygromycin (30μg/ml, final) neomycin (800μg/ml, final) or puromycin (1μg/ml, final). ZC3H4 cDNA (NM_015168.2) was synthesised by Genscript in a pcDNA3.1(+)-C-eGFP vector. The MS2hp-IRES-GFP reporter was made by inserting a BamH1/EcoRV restriction fragment from pSL-MS2-6x (#27118, Addgene) into pcDNA3.1(+)IRES GFP (#51406, Addgene) also digested with BamH1/EcoRV.

### Turbo sample preparation

10 cm dishes at ~80% confluency were labelled with 500 μM biotin for 2 mins and the labelling reaction quenched immediately by washing cells in ice cold PBS. Cells were lysed in RIPA buffer (150 mM NaCl, 1% NP40, 0.5% sodium deoxycholate, 0.1% SDS, 50 mM Tris-HCl at pH 8, 5 mM EDTA at pH 8) containing protease inhibitors (cOmplete mini EDTA-free tablets, Roche) for 30 mins on ice, then clarified *via* centrifugation. 350 uL of washed Streptavidin Sepharose High Performance slurry (GE Healthcare) was incubated in biotinylated or control lysates with inversion at room temperature for 1 hour. Samples were then washed twice with RIPA buffer, twice with Urea buffer (2 M urea, 50 mM Tris HCl pH 8), twice with 100 mM sodium carbonate and once with Tris (20 mM Tris HCl pH 8, 2 mM CaCl_2_). Residual final wash buffer was used to re-suspend the beads, which were then flash frozen in liquid nitrogen and sent for Tandem Mass Spectrometry at University of Bristol Proteomics Facility.

### Identifying mass spectrometry candidates

First, contaminant proteins (i.e keratin) or those that are known to be preferentially biotinylated in ligase assays (i.e. AHNAK) were excluded. Samples with an average abundance ratio of ≤ 0.70 were classified as having a decreased interaction with RNA polymerase II following CPSF30 depletion. Finally, proteins with ≤ 5 peptides were discarded. Remaining candidates were plotted in Figure 1F.

### qRT-PCR

1 μg of total RNA (DNase treated) was reversed transcribed using random hexamers according to manufacturer’s instructions (Protoscript II, NEB); cDNA diluted to 50uL. qPCR was performed using LUNA SYBR (NEB) on a Rotorgene (Qiagen). Fold changes were calculated using Qiagen’s comparative analysis software, graphs were plotted using Prism (GraphPad).

### Antibodies

CPSF30 (A301-585A-T, Bethyl), RNA Pol II (ab817, Abcam), PNUTS (A300-439A-T, Bethyl), WDR82 (D2I3B, Cell Signalling), EXOSC10 (sc-374595, Santa Cruz), ZC3H4 (HPA040934, Atlas Antibodies), HA tag (clone 3f10, 11867423001, Roche), GFP (PABG1, Chromotek), TCF4/TCF7L2 (C48H11, Cell Signalling).

### GFP Trap

10 cm dishes were transfected seeded overnight, washed with PBS, lysed for 30 mins on ice in 1 mL lysis buffer (150 mM NaCl, 2.5 mM MgCl_2_, 20 mM Tris HCl pH 7.5, 1 % Triton X-100, 250 units Benzonase [Sigma]), samples were then clarified through centrifugation (12000xg, 10 mins), then, samples split in two, incubated with 25ul of GFP TRAP magnetic agarose (Chromotek) for 1hr with rotation at 4°C. Beads were washed 5x with lysis buffer and samples eluted by boiling in 2xSDS buffer before analysis by western blotting.

### Nuclear RNA-Seq

Nuclei were extracted from 1x 30mm dish of cells per condition using hypotonic lysis buffer (10 mM Tris pH5.5, 10 mM NaCl, 2.5 mM MgCl_2_, 0.5% NP40) with a 10% sucrose cushion and RNA was isolated using Tri-reagent. Following DNase treatment, RNA was Phenol Chloroform extracted and ethanol precipitated. After assaying quality control using a Tapestation (Agilent), 1 μg RNA was rRNA-depleted using Ribo-Zero Gold rRNA removal kit (Illumina) then cleaned and purified using RNAClean XP Beads (Beckman Coulter). Libraries were prepared using TruSeq Stranded Total RNA Library Prep Kit (Illumina) and purified using Ampure XP beads (Beckman Coulter). A final Tapestation D100 screen was used to determine cDNA fragment size and concentration before pooling and sequencing using Hiseq2500 (Illumina).

### ChIP-Seq

ChIP libraries were prepared using SimpleChIP® Plus Enzymatic Chromatin kit (9005, Cell Signalling) according to manufacturer’s instructions. 5μg of RNA Pol II (abcam, 8WG16) or ZC3H4 (HPA040934, Atlas Antibodies) were used for immunoprecipitation, Dynabeads α-Mouse / α-Rabbit (Life Technologies) were used for capture.

### Chromatin RNA isolation

HCT116 cells were scraped into PBS, pelleted, incubated in hypotonic lysis buffer (HLB; 10 mM Tris.HCl at pH 7.5, 10 mM NaCl, 2.5 mM MgCl2, 0.5% NP40), underlayered with 10% sucrose (*w/v* in HLB) on ice for 5 mins, then spun at 500 xg to isolate nuclei. Supernatant and sucrose was removed and nuclei re-suspended in 100 μL of NUN1 (20 mM Tris-HCl at pH 7.9, 75 mM NaCl, 0.5 mM EDTA, 50% glycerol, 0.85 mM DTT), before being incubated with 1 mL NUN2 (20 mM HEPES at pH 7.6, 1 mM DTT, 7.5 mM MgCl_2_, 0.2 mM EDTA. 0.3 M NaCl, 1 M urea, 1% NP40) on ice for 15 mins. Samples were spun at 13, 000 xg to pellet chromatin, this was dissolved in Trizol and RNA extracted.

### Colony formation assays

*ZC3H4-DHFR* cells were seeded into 100mm dishes and maintained in the presence or absence of TMP for 10 days, with media replaced every 3 days. Colonies were fixed in ice cold methanol for 10 mins and then stained with 0.5% crystal violet (in 25% methanol) for 10 minutes.

### XRNAX

We essentially followed the protocol of (Trendel et al., 2019). HCT116 cells were grown overnight in the presence or absence of doxycycline in 10 cm dishes. 24 hr later, dishes were washed with PBS, UV cross linked (Stratalinker 1800 150 mJ/cm2), or not, then re-suspended in 4.5 mL Trizol (Sigma). 300 uL of chloroform was added, samples agitated on a ThermoMixer (Eppendorf) for 5 mins, span at 12000xg for 15 minutes, then the interphase carefully aspirated into fresh tubes. The interphase was washed thrice with Tris-SDS (10 mM Tris-HCL pH 7.5, 1 mM EDTA, 0.1 % SDS), before being dissolved in 1 mL Tris-SDS. 1 uL glycogen, 60 uL of 5M NaCl and 1 mL isopropanol were added and samples precipitated at −20’C for 10 minutes, then pelleted at 18, 000xg for 15 mins. Precipitated protein was washed with 70 % ethanol, air dried, re-suspended in 180 uL water and pellets dissolved on ice. DNA was removed via TurboDNAse (ThermoFisher) treatment, before samples were re-pelleted, re-dissolved in RNase buffer (150 mM NaCl, , 20 mM Tris-HCL pH 7.5, 2.5 mM MgCl2) and RNA digested with RNase H (NEB) and 1 uL of RNAse T1 (Roche). 4 x SDS loading buffer was added before gel electrophoresis and western blotting.

### Computational analysis

All sequencing data was uploaded to the Galaxy web platform and processed as detailed below; usegalaxy.org and usegalaxy.eu servers were used.

### Datasets (GEO accessions)

Data newly generated in this paper (GSE163015); Pol II HEPG2 ChIP-seq (GSE32883); ZC3H4 HEPG2 ChIP-seq (GSE104247); DIS3-AID HCT116 RNA-seq

(GSE120574); INTS1 RNAi Chromatin-associated RNA-seq (GSE150238).

### RNA-Seq alignment

FASTA files were trimmed using Trim Galore! and mapped to GRCh38 using HISAT2 using default parameters (Kim et al., 2015). Reads with a MAPQ score of ≤ 20 were removed from alignment files using SAMtools (Li et al., 2009). Finally, BigWig files were generated using DeepTools and visualised using IGV (Ramirez et al., 2014).

### ChIP alignment and visualisation

All samples were mapped against GRCh38 using BWA, default settings. Reads with a MAPQ score of ≤ 20 were removed along with PCR duplicates from alignment files using SAMtools. Processed BAM files were converted to BigWig files using DeepTools: all samples were normalised to RPKM with a bin size of 1. Aligned files were visualised using IGV.

### ChIP peak calling

For ZC3H4 and INPUT, broad peaks were called separately using MACS2 with a changed “lower mfold” (2) and default settings. For each experiment, bedtools was used to establish common peaks from individual reps (Intersect Intervals), creating a bed file of high confidence peaks – for ZC3H4, peaks called in INPUT sample were subtracted *via* bedtools. All bed files were annotated and plotted in R using ChipSeeker (Yu et al., 2015).

### Gene heatmaps

For ChIP heatmaps, computematrix (DeepTools) was used to generate score files from ChIP bigwig files using an hg38 bed file; parameters used for each heat map are detailed in figure legends. Plots were redrawn in R.

### Super-enhancer metaplots

A bed file with the coordinates of super-enhancer locations from dbSUPER in HCT116 cells was used as a basis (Khan and Zhang, 2016). All regions that had clusters of MED1, Pol II and H3K27ac ChIP signal were retained as *bone fide* regions of interest, those without were discarded. A log_2_ ratio of experiment vs input was prepared using BamCompare of DeepTools - for RNAseq metaplots, BAM files were split by strand. A score file for the regions in the amended SE bed file was generated *via* the computematrix function of DeepTools using the log_2_ BamCompare output file. Results were plotted in R-studio using ggplot2.

### Gene plots and metaplots

Read-through metaplots were generated as described (Davidson et al., 2019). For identifying ZC3H4 PROMPT regions ncRNA genes were filtered from hg38 refgene gtf file to give protein-coding genes that were used with feature counts on siCont RNAseq (Liao et al., 2014), to gain read count and gene length. Transcripts per million (TPM) were calculated for each gene and genes with an expression of < 5 were filtered out to give a list of expressed genes. Next, divergent promoters, or genes with neighbours within 5kb of their promoter were excluded to minimise background. Finally, this gene list was converted to a bed file, then computematrix (DeepTools) used to generate a score file from log_2_ Scramble Vs Experiment bigwigs; results were plotted in R.

### ZC3H4 homolog identification

To identify ZC3H4 homolog protein sequences, sequences from UniRef100 (UniProt Consortium, 2014) were searched using a profile HMM search: ‘hmmsearch’, part of HMMer V3.2.1 (Eddy, 2011), with PANTHER (Mi et al., 2019) hidden Markov model PTHR13119, corresponding to zinc finger CCCH-domain containing proteins. Profile HMM search hits were filtered using a 1e-100 e-value threshold; this search identified 1513 UniRef100 sequences with PTHR13119 domains (representing a total of 1646 UniProtKB sequences).

### Phylogenetic tree reconstruction

Identified PTHR13119 domains were aligned using MAFFT v7.4 (Katoh and Standley, 2013); sites composed of more than 75% of gaps were removed from the multiple sequence alignment with trimAl (Capella-Gutierrez et al., 2009). The PTHR13119 domain phylogeny was reconstructed under maximum likelihood with IQ-TREE v1.6.9 (Nguyen et al., 2015). The best-fitting substitution matrix was determined by ModelFinder (Kalyaanamoorthy et al., 2017), as implemented in IQ-TREE. Branch support values were based on 1000 ultrafast bootstraps (Minh et al., 2013). Phylogenetic Tree figure was rendered with iToL (Letunic and Bork, 2019).

### Primers, siRNAs and other nucleic acid sequences

Sequences provided in Supplementary Excel file 4.

---

**Figure S1.**
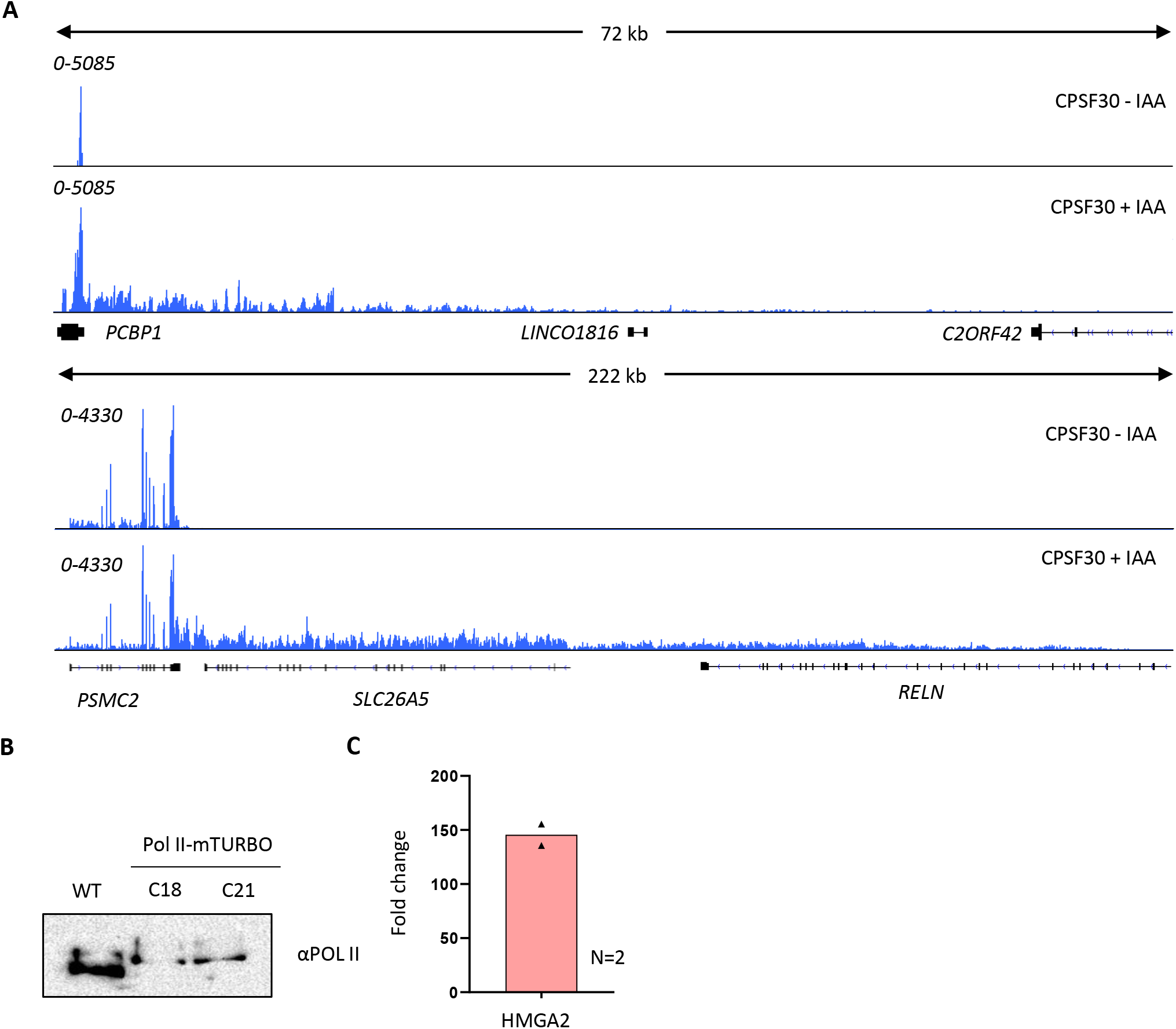
a) IGV track view of the transcription termination defect at *PCBP1* and *PSMC2* genes in the presence (CPSF30-IAA) or absence of CPSF30 (+IAA). Signal is RPKM. b) Western blot demonstrating bi-allelic modification of RPB1 (Pol II) with mTurbo. Two clones (C18 and C21) are shown against cells (WT) not modified at RBP1. The upshift of Pol II signal shows bi-allelic modification of RPB1. C21 was used for the experiment in figure 1. c) qRT-PCR of total RNA isolated from *CPSF30-mAID: RPB1-mTurbo* cells treated or not with auxin (3hr). An amplicon located ~10kb downstream of the *HMGA2* PAS was used to assay transcriptional read-through presented as a fold change versus minus auxin after normalising to spliced actin mRNA. n=2.

**Figure S2.**
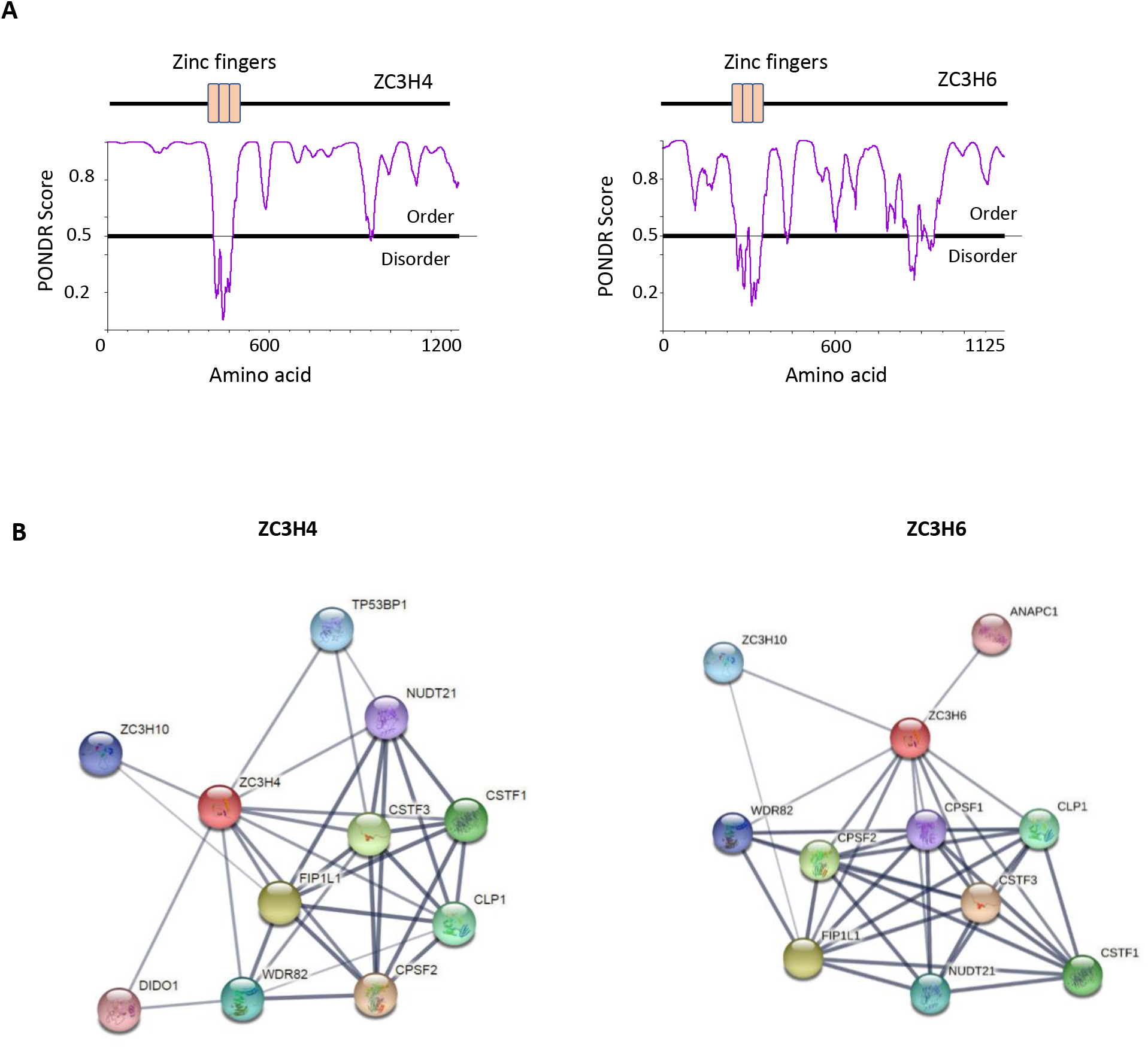
a) Schematic of ZC3H4 and ZC3H6 showing the three CCCH zinc finger domains. A predictor of natural disordered region (PONDR) analysis shows that the only ordered region coincides with these domains. Graph generated via PONDR.com, set to VSL2. b) String analysis of ZC3H4 and ZC3H6 indicate interactors with 3’ end processing complex. Image was taken from string.db.org, confidence value was set to medium (0.4). The thickness of lines between nodes denotes confidence of interaction.

**Figure S3.**
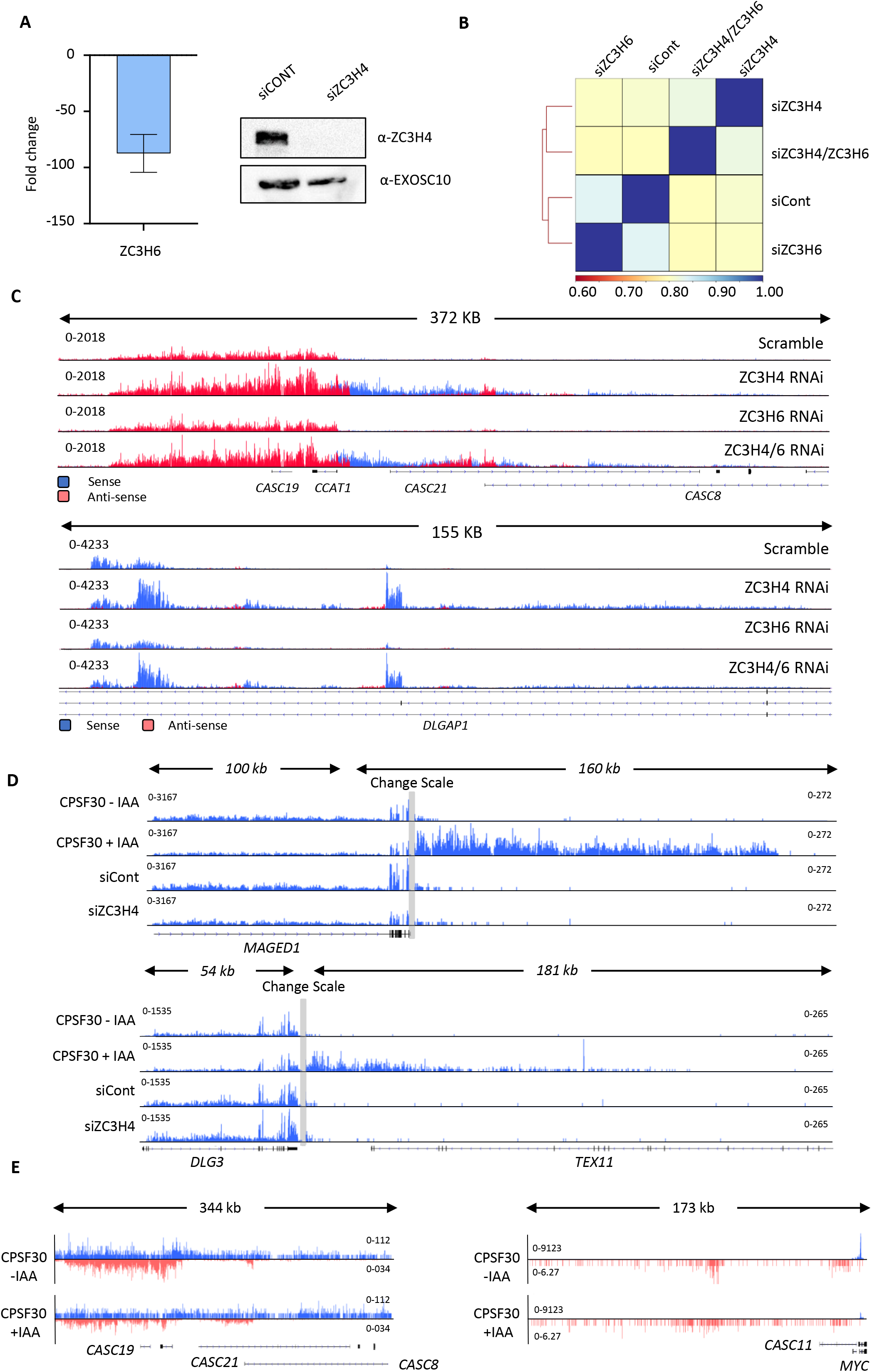
a) qRT-PCR and western blotting evidence of the effectiveness of ZC3H6 and ZC3H4 depletion respectively. Graph shows fold reduction of ZC3H6 mRNA in cells treated with ZC3H6 siRNAs versus those transfected with control siRNA. N=3, error bars are SEM. Western blotting of ZC3H4 in HCT116 cells treated with control siRNAs or ZC3H4-targeting siRNAs. The blot was probed with a ZC3H4 antibody revealing strong depletion versus the EXOSC10 loading control. b) Pearson’s correlation of siControl, siZC3H4, siZC3H6 and siZC3H4+6 of RNAseq BAM files performed by DEEPTOOLS. c) IGV traces exemplifying two genomic regions with clear RNA accumulation following ZC3H4 depletion. While this is also seen following ZC3H4/ZC3H6 co-depletion, it is not evident following loss of ZC3H6 alone. This was generally seen, supporting the correlation analysis in b). y-axis scale is RPKM. d) IGV tracks of individual genes (*MAGED1* and *DLG3)* to exemplify the lack of 3’ termination defect following ZC3H4 depletion. As a control for *bone fide* read-through, the same tracks are shown in samples obtained from *CPSF30-mAID* cells treated or not (3hr) with auxin. Grey bar indicates a change in scale (left scale for upstream of PAS; right for downstream) so that comparatively weaker read through signals can be visualised next to the gene body. y-axis scale is RPKM. e) CPSF30 depletion shows little effect at super-enhancers and PROMPTs. IGV track view of the *MYC* super-enhancer and PROMPT in RNA-seq data obtained from *CPSF30-mAID* cells treated or not (3hr) with auxin. y-axis scale is RPKM.

**Figure S4.**
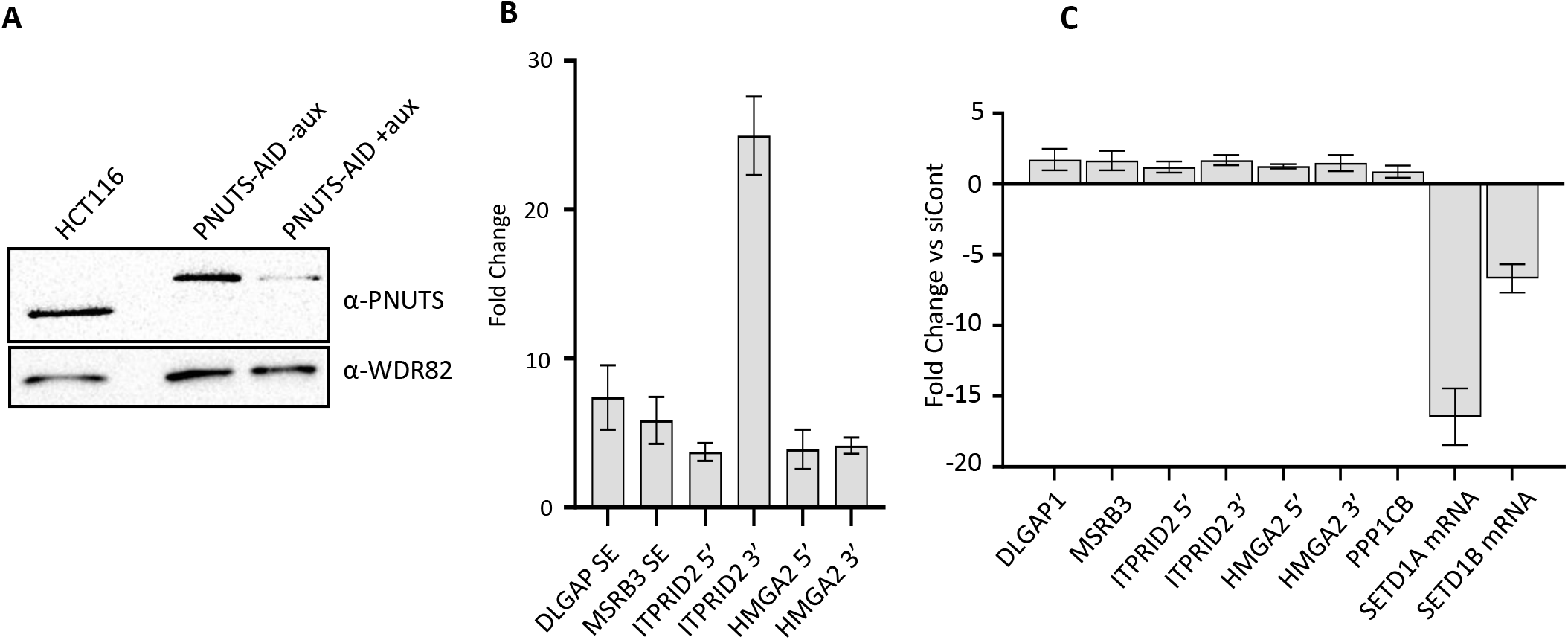
a) Western blot showing acute depletion of PNUTS, tagged with an auxin-inducible degron (PNUTS-AID). Blot shows extracts from unmodified HCT116 cells and *PNUTS-AID* cells which were treated or not with auxin (3hr). WDR82 is used as a loading control. b) qRT-PCR of PROMPT and SE transcripts in *PNUTS-AID* cells treated or not with auxin (3 hr). Graph shows fold change by comparison with non-auxin treated cells following normalisation to spliced actin transcripts. N=3. Error bars are SEM. c) qRT-PCR analysis of PROMPT and SE transcripts in HCT116 cells treated with control siRNAs or siRNAs against both SETD1A and B. Quantitation shows fold change versus cells transfected with control siRNAs following normalisation to spliced actin levels. Note that the PROMPT and SE targets that are increased following ZC3H4 loss show relatively little change following depletion of SETD1A and B. The success of the RNAi is indicated by the strong reduction of SETD1A and B mRNAs. N=3. Error bars show SEM.

**Figure S5.**
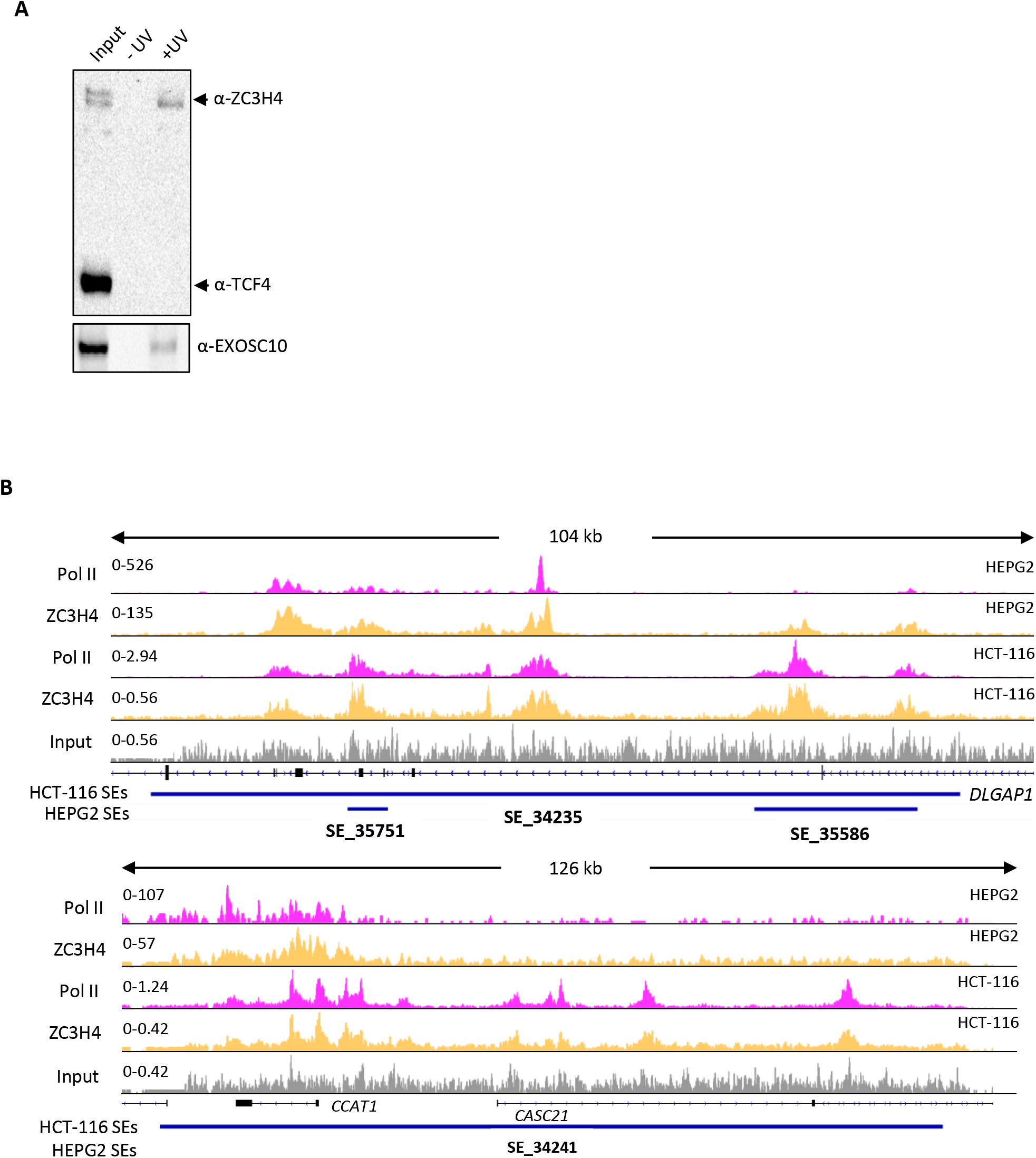
a) XRNAX analysis of ZC3H4 RNA binding in cells. Samples show input and those isolated following UV treatment or not. Bands representing each protein are labelled accordingly. ZC3H4 is recovered in a UV-dependent manner indicating that it is directly bound to RNA in cells. The same is true of EXOSC10 that, as an exoribonuclease, acts as a positive control. TCF4 is a DNA binding transcription factor and acts as a negative control. b) ZC3H4 only marks transcribed super-enhancers. IGV track view of ZC3H4 and Pol II occupancy at two different super-enhancers, *DLAGAP1* present in both HEPG2 and HCT116 (top tracks) and the MYC super-enhancer (bottom tracks) that is only present in HCT116 cells. HEPG2 and HCT116 super-enhancer annotation is under each track as blue bars and was obtained from dbSUPER. Y-axis shows CPM.

